# Rare earth elements (REE) in freshwater, marine, and terrestrial ecosystems in the eastern Canadian Arctic

**DOI:** 10.1101/174870

**Authors:** Gwyneth Anne MacMillan, John Chételat, Joel Heath, Raymond Mickpegak, Marc Amyot

**Affiliations:** Centre for Northern Studies, Department of Biological Sciences, University of Montreal, Montreal, QC, Canada, H2V 2S9; Environment and Climate Change Canada, National Wildlife Research Centre, Ottawa, ON, Canada, K1A 0H3; Arctic Eider Society, St. John’s, NL, Canada, A1C 3Z6; Sakkuq Landholding Corporation, Kuujjuaraapik, QC, Canada, J0M 1G0

**Keywords:** Metals, Rare Earth Elements (REE), Lanthanide, Arctic, Subarctic, Bioaccumulation, Stable Isotope

## Abstract

Few ecotoxicological studies exist for rare earth elements (REEs), particularly field-based studies on their bioaccumulation and food web dynamics. REE mining has led to significant environment impacts in several countries (China, Brazil, U.S.), yet little is known about the fate and transport of these contaminants of emerging concern. To understand how REEs behave in pristine northern food webs, we measured REE concentrations and carbon and nitrogen stable isotope ratios (∂^15^N, ∂^13^C) in biota from marine, freshwater, and terrestrial ecosystems of the eastern Canadian Arctic (N=339). Northern ecosystems are potentially vulnerable to REE enrichment from prospective mining projects at high latitudes. Wildlife harvesting and tissue sampling was partly conducted by local hunters through a community-based monitoring project. Results show that REE generally follow a coherent bioaccumulation pattern for sample tissues, with some anomalies for redox-sensitive elements (Ce, Eu). Highest REE concentrations were found at low trophic levels, especially in vegetation and aquatic invertebrates. Terrestrial herbivores, ringed seal, and fish had low REE levels in muscle tissue (<0.1 nmolg^-1^), yet accumulation was an order of magnitude higher in all liver tissues. Age- and length-dependent REE accumulation also suggest that REE uptake is faster than elimination for some species. Overall, REE bioaccumulation patterns appear to be species- and tissue-species, with limited potential for biomagnification. This study provides novel ecotoxicological data on the behaviour of REE in ecosystems and will be useful for environmental impact assessment of REE enrichment in northern regions.

## INTRODUCTION

Rare earth elements (REEs) are a chemically-similar group of emerging contaminants, which includes the 15 trivalent lanthanide metals, as well as scandium (Sc) and yttrium (Y). Not particularly rare, REEs are increasingly exploited for critical uses in high-tech industries, including electronics, medicine, clean energy, and agriculture.^1, 2^ REEs are used for magnets, metal alloys, catalysts, fertilisers and ceramics, as well as for eutrophication management systems in freshwaters.^3, 4^ Increasing emissions have led to significant release of REEs into the environment, yet knowledge of their fate and impact on natural ecosystems is limited. Most existing studies use REEs to trace geochemical processes in natural systems and very few studies have examined the ecotoxicology and/or environmental impacts of these metals. The majority of existing REE ecotoxicity studies are laboratory-based, whereas field measurements of natural background levels, environmental behaviour and bioaccumulation potential in food webs are extremely rare.^5, 6^

Mining and processing of rare earth ore is known to have major environmental impacts, including the production of atmospheric pollution, acidic wastewater, and radioactive tailings.^7–9^ Significant REE enrichment in water, soil and vegetation near mining sites in China has also led to recent concerns about the environmental impacts of REEs themselves.^10–13^ Rivers located near rare earth mines in China, for example, have dissolved REE concentrations three orders of magnitude higher than unperturbed rivers.^8^ A lack of key ecotoxicological data renders environment impact assessment of REEs difficult and hinders the creation of environmental guidelines or thresholds.^2^ REEs were traditionally considered as low risk to environmental or human health because they are lithophilic, hence largely insoluble and immobile (i.e. not bioavailable) in soils.^14, 15^ However, recent laboratory studies show the potential for both toxicity and bioaccumulation of REEs in many species, including microorganisms and phytoplankton,^16^–18 aquatic plants,^5, 19^ terrestrial plants,^20–24^ terrestrial and aquatic invertebrates, ^1, 10, 13, 25^ as well as in fish.^26–28^ Anthropogenic REE enrichment, and subsequent bioaccumulation in biota, may therefore lead to REE concentrations closer to estimated toxicity thresholds, as is the case in the Yellow River region in China.^8^

Rising demand for REEs, and decreasing export from China, have recently led to many new REE mining ventures around the world.^7^ Although no REE production or refining currently occurs in Canada, more than 200 exploration projects are under development, including 11 in an advanced stage. The majority of these projects are found in northern Canada, including five projects in northern Quebec and 2 in the Northwest Territories.^4^ Accelerated warming combined with significant pressure to exploit natural resources mean that Arctic ecosystems are vulnerable to rapid industrial and environmental change. Climate change is accelerated at high latitudes and has been shown to affect contaminant cycling, including for metals.^29^ It is therefore important to consider the environmental impacts of REE enrichment at high latitudes, yet only limited data are available for REE environmental behaviour in northern ecosystems.

The aim of this field-based study was to evaluate the potential for bioaccumulation and trophic transfer of REEs in freshwater, marine, and terrestrial food webs of the eastern Canadian Arctic. Biological samples were collected in 2012, 2014 and 2015 from marine, terrestrial, and freshwater ecosystems near Kuujjuarapik-Whapmagoostui (Nunavik, Quebec). Wildlife harvesting and tissue sampling were conducted by local hunters through a community-based monitoring project. The objectives of this study were a) to assess REE levels in biota of different trophic levels found in freshwater, marine and terrestrial ecosystems in the Arctic, focusing on taxa of importance to Northerners, b) to evaluate the species- and tissue-specific bioaccumulation of REEs in key taxa and c) to trace the trophic transfer of REEs within northern ecosystems using carbon and nitrogen stable isotope measurements of food web structure. Quantifying and tracing REE behaviour in these ecosystems will allow us to better evaluate the potential environmental impact of REE enrichment in the northern environment.

## MATERIALS AND METHODS

### Study sites

Marine, freshwater (lakes and river) and terrestrial ecosystems were sampled near Kuujjuarapik-Whapmagoostui (K-W), a community located in the subarctic taiga in Nunavik, Quebec, Canada (55° 16’ 30’’ N, 77° 45’ 30’’ W). This region encompasses marine, freshwater and terrestrial ecosystems of importance to Northerners, including southeastern Hudson Bay, the Great Whale River, and numerous lakes underlain by discontinuous, scattered permafrost.^30^ All samples were collected within a relatively restricted geographic radius (< 70 km) around K-W. No documented rare earth element deposits or exploitation activities currently exist in this region and our study provides baseline environmental data for REE in the eastern Canadian Arctic.

### Lake sampling

In 2012, replicate samples of benthic invertebrates, zooplankton, and fish were collected from 8 lakes for the analysis of REE concentrations and stable isotope ratios of nitrogen and carbon (∂15N, ∂13C). Bulk zooplankton were sampled by horizontal surface hauls with a large net (1 m diameter, 200 µm mesh). Benthic invertebrates were sampled along the shoreline with a kick net (500 µm mesh), or an Ekman grab for deeper water, and were live-sorted into broad taxonomic groups without depuration. Brook trout (*Salvelinus fontinalis*), the only large-bodied fish species at these sites, were captured with a gill net from 5 of the 8 lakes. Ancillary data collected for the brook trout included total length, fork length, mass, sex and age (estimated by annuli counting of otoliths using the crack and burn method). Surface water and surface sediment (top 1-2 cm, by Eckman grab) were also collected for REE analysis from an inflatable raft with an electric motor. Ultra-trace metal sampling techniques were used to quantify REE concentrations.

### Marine, river, and terrestrial sampling

In 2014 and 2015, wildlife harvesting was organized by the Sakkuq Landholding Corporation through community-based projects on metal bioaccumulation. Local hunters collected tissue samples from biota in Hudson Bay, the Great Whale River and terrestrial ecosystems near K-W. Species were chosen to reflect taxa of importance to local communities as well as a variety of representative trophic levels and ecosystems. The skills, experience and traditional knowledge of hunters were critical for the animal collections, particularly related to the distribution and seasonal movements of local populations. Record sheets were used to record data on the location, size, and sex of harvested animals. Ancillary data collected for ringed seals included length, blubber thickness, and axial and maximum girth.

Marine sampling included tissue collection (muscle, liver) of a top marine predator, the ringed seal (*Phoca hispida*), as well as a benthic molluscivore, the common eider (*Somateria mollissima*). Marine invertebrates were also collected from coastal sites: a planktonic feeder, the blue mussel (*Mytilus edulis*, all tissues without shell) and a benthic feeder, the sea urchin (order Echinoida, gonads). For the river samples, anadromous freshwater fish were collected from the mouth of the Great Whale River, including brook trout (*Salvelinus fontinalis*) and lake whitefish (*Coregonus clupeaformis*). Terrestrial sampling included tissue collection (muscle, liver) from three terrestrial herbivores: snowshoe hare (*Lepus americanus*), willow ptarmigan (*Lagopus lagopus*), and caribou (*Rangifer tarandus*). Sampled terrestrial vegetation included above-ground tissues (stems and leaves) from 1) vascular plants: crowberry (*Empetrum nigrum*), Labrador tea (*Rhododendron* sp.), and bearberry (*Arctostaphylos alpina*), and 2) non-vascular plants: lichens (fruticose type, ground and tree) and moss (*Spagnum* sp.). The surfaces of plant samples (e.g., leaves) were not cleaned with ultra-pure water and therefore, REE measurements represent both internal accumulation and external adsorption on surfaces, which is relevant for estimating metal exposure to grazing herbivores.

### Rare earth element analysis

Sediment and biological samples were stored at −20°C, freeze-dried, and homogenized before analysis for REEs by inductively coupled plasma mass spectrometry (ICP-MS, Perkin-Elmer NexION 300x) following microwave digestion. From 0.07 - 0.20 g (median 0.10 g) of sample was weighed into pre-washed Teflon tubes (HNO_3_ 45%, HCl 5%) and digested with 3 mL of trace metal grade HNO_3_ (70%) for 15 minutes at 170°C. Two more 15 minute cycles were completed after adding 0.5 - 1.0 mL of OPTIMA grade hydrogen peroxide (30% H_2_O_2_) before each cycle. Digested samples were diluted with ultra-pure water (MilliQ, 18.2 MΩ•cm) to a volume of 50 mL and then re-diluted (1:2) into trace metal clean falcon tubes. Surface water samples were filtered on a clean Teflon filtration tower (HCl 10%) and preserved with HNO_3_ (2%) before analysis. Vegetation samples were digested using longer cycles (30 min) with more H_2_o_2_ (1.0 mL each cycle) to permit a more complete digestion of tough plant material. Samples with low biomass (benthic invertebrates, 0.005 - 0.015g) were digested over a longer period at a lower temperature (room temperature for 24h, hot plate for 2h at 80°C) using equivalent volumes of HNO_3_ (70%) and H_2_O_2_ and diluted to 10 mL with ultra-pure water.

ICP-MS detection limits were sufficiently low (0.0001 - 0.0022 nmolg^-1^) to quantify samples with low REE concentrations (Table S1). REE digestions included analytical blanks and appropriate reference standards, including sediment (STSD-1, stream sediment, CCRMP, CANMET), animal tissues (BCR 668 mussel tissue, IRMM) and plants (BCR 670 aquatic plant, IRMM). Average uncertainly of the method was ± 10% for samples with high concentrations and ± 40% for samples with low concentrations close to detection limits. Average (min-max) recovery of reference material was 87% (79 - 101%) for animal tissue, 84% (67 – 117%) for plant tissue, and 70% (40 – 99%) for sediment (Table S2). These values are consistent with average recoveries reported in the literature.^5, 27, 31^ It is important to note that certified values are classified as “total” concentrations and use multi-acid dissolution (HNO_3_, HF). Some of the variability in analytical recovery could be explained by the less aggressive or “partial” extraction methods used in this study to estimate the labile, bioavailable REE concentration in samples.^32^

### Stable Isotope Analysis

Dry sediment and biological samples were weighed into tin capsules and analyzed for stable isotope ratios (δ^13^C, δ^15^N) using an elemental analyser interfaced with an isotope ratio mass spectrometer (IR-MS, Thermo Delta Advantage) at the G.G. Hatch Lab (U. of Ottawa). Only δ^15^N values were analyzed for sediments because samples were not acidified to remove carbonates. Stable isotope ratios are reported in Delta (δ) notation, the unites are parts per thousand (‰) and defined as δX= ((Rsample-Rstandard))/Rstandard) x 1000 where X is ^15^N or ^13^C, and R is the ratio of the abundance of the heavy to the light isotope. For freshwater lakes, nitrogen stable isotope ratios (δ^15^N) were adjusted for among lake differences in baseline values using δ^15^N values from lake sediment. The adjusted value (δ^15^N_adj_) allows for comparison between lakes by accounting for variation in baseline δ^15^N across different ecosystems. Quality assurance included triplicate analyses of an internal standard (analytical precision of 0.2 ‰) and duplicate analyses of 10% of samples.

### Data handling

In this study, 15 of the 17 rare earth elements were used to calculate the sum of all detected REE (∑REE) in nmol/g (biota and sediments) or nmol/mL (water). Promethium (Pm) was not included as it does not occur naturally and results for scandium (Sc) were excluded due to analytical interference. Previous studies have found that Sc (^45^Sc^+^) had false high readings due to interference with sample organic content^33, 34^ and that Sc is not strongly correlated with other REE.^27^ Detection frequencies were variable for the 15 REEs and different taxonomic groups in this study. Non-detected elements were often from the HREE group, except for Y which was the most widely-detected element (90% of samples). Y, La, and Nd were detected in over 80% of samples, whereas Tm and Lu were detected in < 50% of samples. 100% of elements were detected in the aquatic invertebrate and plant samples, but low detection frequencies (< 30% of elements) were found in samples from aquatic vertebrates (seal, eider fish) (Table S3).

Geometric means (the antilog of the mean of the logarithmic values of the data set) were used to calculate average ∑REE within taxonomic groups to measure central tendency with high intra-group variation. Analytical blank values were subtracted from the sample values for each element. Sample measurements of REEs below detection limits (<DL) were estimated as the concentration of half the detection limit value, except when applying normalisation as in Fig. 4. Different species were pooled together within ecosystems to compare taxonomic groups. For example, terrestrial vegetation includes vascular plants and non-vascular plants, and freshwater benthic invertebrates include amphipods, caddisflies, anisoptera, and corixidae. All tissue concentrations are presented on a dry-weight basis.

For data analysis, we divided REEs into two groups based on physicochemical parameters, the light REEs (LREE) from La-Gd and the heavy REEs (HREE) from Tb-Lu and Y. Other authors divide REE into three groups and the specific elements placed in each group vary between studies.^23, 27, 35^ To graphically compare REE abundances from different samples, individual element concentrations were normalised based on their geological abundance using a standard (Post Archean Australian Shale or PAAS).^36^ Normalisation eliminates the Oddo-Harkins effect (or saw-tooth pattern) and deviations from a horizontal line after normalization indicate natural or anthropogenic enrichments (or depletions) of elements (or anomalies). Eu and Ce anomalies (∂Eu and ∂Ce) were calculated by ∂Eu = Eu_PAAS_ / (Sm_PAAS_ x Gd_PAAS_)^0.5^ and ∂Ce = Ce_PAAS_ / (La_PAAS_ x Pr_PAAS_)^0.5^ where PAAS indicated Post Archean Shale Standard normalized values.^37^

### Statistical analyses

All statistical analysis was performed in R version 3.3.1 (R Core Development Team, 2016). ^38^ Significant levels were *α* < 0.05. Concentration data (muscle, liver) for REEs (individual elements and ∑REE) were log_10_ transformed to reduce skewness and the influence of outliers. Stable isotope, age, and size (length, girth) values followed normal distributions. Correlations between REE concentrations in tissue samples were examined by principal component analysis (PCA) on centered and scaled data (R: vegan package).^39^ Simple linear regression analysis was conducted to assess the relationship between LREE and HREE concentration using all available data (N=339). Comparisons of log_10_-SREE concentrations between different taxonomic groups and tissue types were conducted using Welch’s analysis of variance (ANOVA) and Games-Howell post-hoc tests; this is a nonparametric approach that does not assume equal variance or sample size between groups (R: *userfriendlyscience* package).

Linear mixed effects model analyses (LMM) were performed on brook trout (lakes only, N=58) and ringed seal (N=23) datasets using R package *lme4*.^40^ This approach evaluated whether tissue REE concentration varied with animal size and sex, while controlling for habitat (lake ID) and year collected. Only liver concentrations were used in the models as muscle concentrations were close to detection limits. Variables were standardized (centered, reduced) and were excluded from models when highly collinear (Pearson’s product-moment correlation coefficients > 0.8). Random intercepts, slopes and interaction terms were tested and removed when removal improved (or did not significantly change) model fit using the Akaike information criterion (AICc, R: *AICcmodavg* package). A brook trout outlier was excluded to improve model fit. Marginal R^2^ (variance explained by fixed factors) and conditional R^2^ (variance explained by fixed and random effects) were obtained from the models fitted through restricted maximum likelihood analysis.^41^ Model validation for all linear models was performed by visual inspection of residual plots, which did not show deviation from homoscedasticity or normality.

## RESULTS AND DISCUSSION

### REE behaviour in northern ecosystems

Concentration patterns show that all 15 REEs are highly correlated with each other in tissue samples. Simple linear regression analysis of LREE versus HREE concentrations (log-scaled nmolg^-1^) in biota showed that concentrations of heavy and light elements were strongly and positively correlated in tissues (Fig 1). Regression coefficients were significant for vertebrate muscle and liver tissues (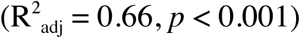) and for plants and invertebrates (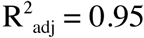, *p* < 0.001). Almost all individual samples were found below the 1:1 slope line (dotted line) on the scatterplot, demonstrating that LREE were consistently more concentrated than HREE in tissues (Fig. 1). On average, LREE comprised 86 ± 19% (mean ± SD) of the total REE content in biota.

**FIGURE 1:**
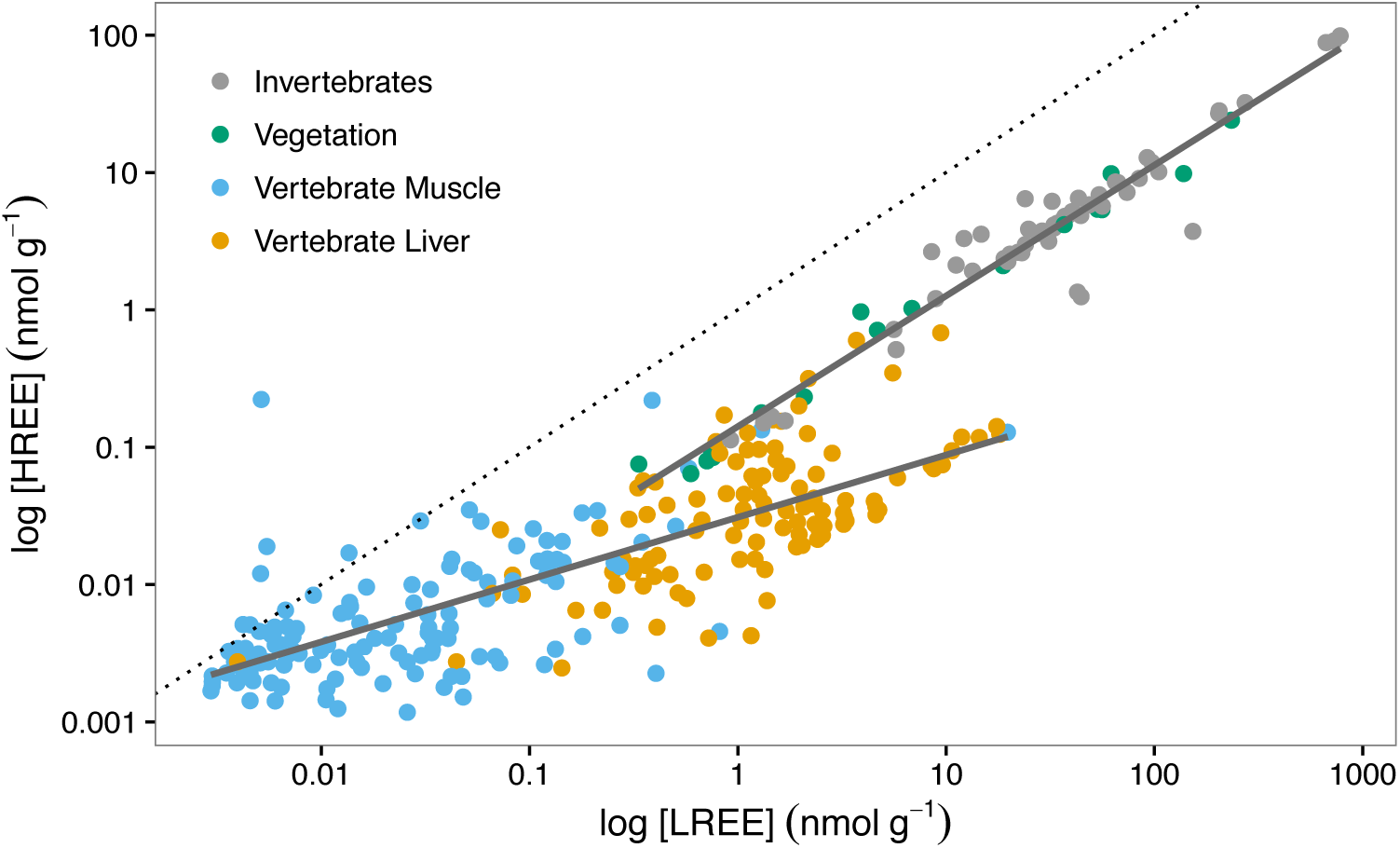
Relationship between LREE and HREE concentrations (log-scaled nmolg^-1^) in biota (N=339) from all ecosystems (vertebrate muscle and liver: 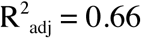, *p* <0.001 invertebrates and vegetation: 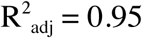, *p* <0.001). Dotted line shows 1:1 slope. Dots show individual samples of invertebrates (marine, freshwater), vertebrate liver and muscle (marine, freshwater, terrestrial), and vegetation (terrestrial: vascular plants, moss, lichen).

REEs are known to be a strongly coherent and predictable group of elements in surface waters, soils and rocks based on chemical similarities (trivalent, electropositive, insoluble) and geochemical behaviour. However, information is scarce on REE behaviour when undergoing bioaccumulation in living organisms. Most previous studies have focused only on bioaccumulation patterns 3 or 4 elements (mainly LREE).^2^ By examining all 15 REEs, we have identified a relatively uniform trend of REE bioaccumulation among a wide variety of taxonomic groups in northern ecosystems. Strong covariance between all 15 REE suggests that ∑REE (i.e. the sum of all REEs) can be used as a good proxy for individual REE concentrations in tissues. Different slopes for LREE vs. HREE concentrations between taxonomic groups (Fig. 1) may be due to analytical variability at low concentrations (e.g. in vertebrate muscle) or may indicate finer scale differences in REE bioaccumulation patterns between vertebrate tissues and invertebrates/plants.

REE bioaccumulation in biota typically display a saw-tooth pattern or “REE pattern” following Oddo-Harkins rule.^5^ This pattern is due to a) log-linear decrease in concentration with atomic number and b) higher concentrations in even-numbered elements over adjacent odd-numbered ones. Finding this pattern mirrored in tissues (Fig. S1, left panels) indicates that REE bioaccumulation mirrors environmental REE concentrations, which are strongly conserved in soils, sediments, and water relative to the Earth’s crust. Our results also show that LREE were consistently more concentrated than HREE in tissues (Fig. 1). This difference may be due to naturally lower concentrations of HREE in the environment or to the hypothesis that REE bioavailability decreases with increasing atomic number due to increasing ligand stability in HREE. For example, Weltje et al. proposed that the presence of ligands in natural surface and pore waters decreased HREE bioavailability and hence bioaccumulation in aquatic ecosystems.^5^

### REE bioaccumulation and biomagnification

Little information exists on REE concentrations in biota from natural ecosystems. In this study, mean ∑REE concentrations varied widely from 0.013 to 103 nmolg^-1^ dry weight (geometric mean or GM) (Fig. 2, Table S4). The highest ∑REE concentrations were found in biota at the base of the food web, especially in lichen/moss (42 ± 81), marine invertebrates (sea urchins 17 ± 7.6; blue mussels 38 ± 5.9), and freshwater invertebrates (benthic invertebrates 33 ± 85, zooplankton 103 ± 484) (GM ± SD, nmolg^-1^). Biota at the base of the food web (vegetation, invertebrates) had significantly higher ∑REE concentrations than vertebrate samples (muscle only) from the same ecosystem (Welch’s ANOVA, F = 108.3, p < 0.001). Mean SREE concentrations in lichen and moss samples were more than 35 times more concentrated than in vascular plants collected in the same general area (GM of 41.5 versus 1.12 nmolg^-1^) (Welch’s ANOVA, F = 81.8, p < 0.001) (Fig 2).

**FIGURE 2:**
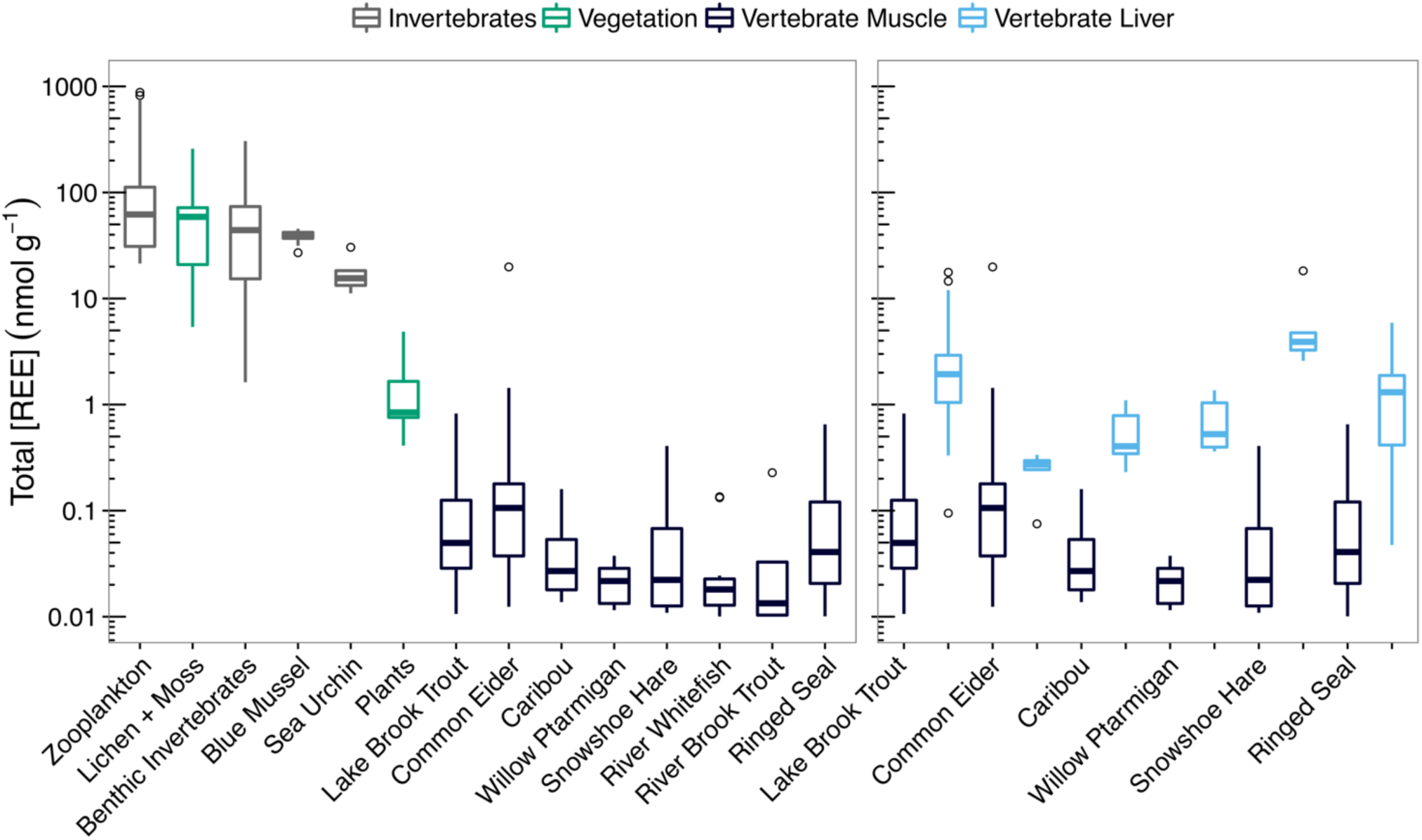
Boxplot of ∑REE concentrations (log-scaled nmolg^-1^) showing median ± SD; dots are outliers (N = 5 - 60 see Table S4). Left panel shows ∑REE concentrations in descending order for each taxonomic group (only vertebrate muscle tissues). Right panel shows vertebrate ∑REE concentrations in muscle compared to liver tissues from the same animals. Differences in mean ∑REE between taxonomic groups were compared using Welch’s ANOVA with Games-Howell post-hoc tests.

Although it is known that REE are bioavailable to lichens and plants,^5, 6, 42^ our results suggest that REE are also widely bioavailable to aquatic invertebrates (both freshwater and marine) and these taxa can therefore be considered good bio-indicators of REE contamination. A handful of marine studies have reported relatively high levels for individual REE in plankton,^43^ flying squid,^44^ and scallops^45^, and only one study has previously measured REE levels in freshwater invertebrates (snails)^5^. In this study, vascular plants had a mean SREE concentration of 1.12 nmolg^-1^ or 0.15 mgkg^-1^ (Table S4), which falls at the lower end of the range of previous reported values for above-ground tissue concentrations (0.06 to 1.6 mgkg^-1^).^46–48^ Plants usually reflect the REE distribution and exchangeable (i.e. soluble) soil concentrations in substrate soils, with higher bioaccumulation in plants on low pH soils.^24^ Previous studies showed that lichens accumulate two-fold higher concentrations of REEs than vascular plants^49^, whereas our results show that natural REE levels can be an order of magnitude higher for lichens and moss. Unlike vascular plants, lichens and moss accumulate elements from atmospheric deposition (rainfall, dust), which may be a pathway for greater REE bioaccumulation than the uptake of REEs by root systems in vascular plants.^47^

Comparisons between REE concentrations and δ^15^N showed that ∑REE decrease with trophic level across marine, freshwater and terrestrial ecosystems (Fig. 3). When compared to lower trophic levels, ∑REE values were roughly an order of magnitude lower in vertebrate liver tissues and 2 orders of magnitude lower in muscle. A decrease in concentration with δ^15^N demonstrates that there is limited potential for biomagnification of REEs in these northern ecosystems. Previous laboratory studies have shown that REEs bioaccumulate but do not appear to biomagnify within aquatic or terrestrial microcosms.^2, 5, 6^ Laboratory studies, however, may not accurately represent REE behaviour within natural ecosystems, because they are simplified systems within a limited range of parameters. To our knowledge, only two studies have measured REE levels in coexisting organisms found at different lower trophic levels within natural food webs.^5, 43^ Our field-based research confirms the limited potential for REE biomagnification within 3 different northern ecosystems.

**FIGURE 3:**
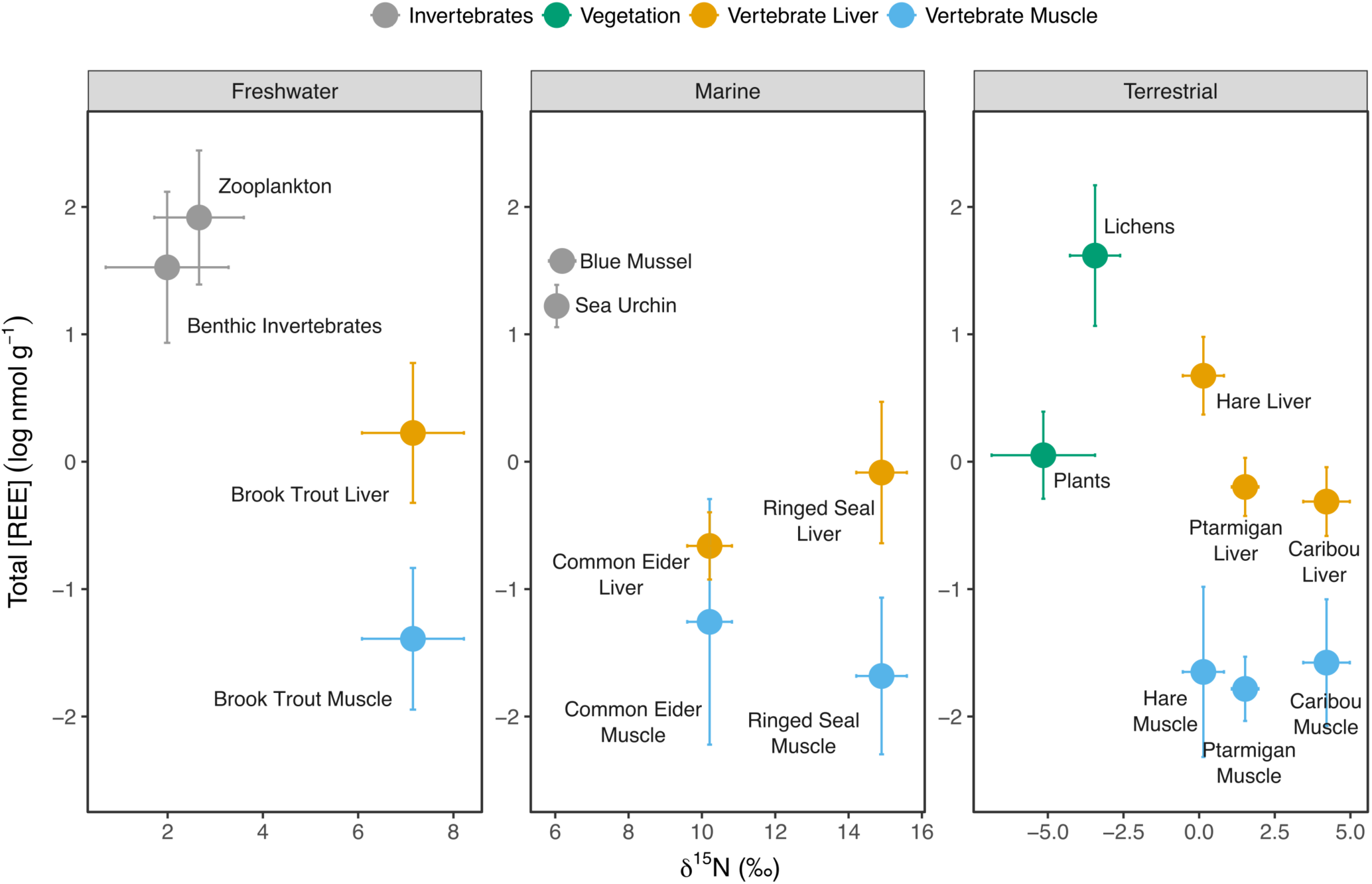
Relationship between mean ∂15N (‰) and mean ∑REE concentrations (log_10_ nmolg^-1^) by taxonomic groups in freshwater (lakes only), marine and terrestrial ecosystems. Values shown are mean ± standard deviation. ∂15N values were adjusted for baseline ∂15N variation in freshwater lakes using ∂15N sediment values (∂15N_adj_). Sample size (N) varies from 5 to 60 (Table S4).

### Tissue-specific REE bioaccumulation

Vertebrates from all ecosystems had REE muscle concentrations that were orders of magnitude lower than ∑REE concentrations in biota near the base of the food web (Fig. 2, right panel). As in this study, previous research has also found that REE are commonly not detected or found at trace levels in fish muscle.^27, 37^ Similarly, common eider, ringed seal, caribou, ptarmigan, and snowshoe hare had low mean REE muscle concentrations, less than 0.1 nmolg^-1^ (or 0.01 mgkg^-1^). To the best of our knowledge, this is the first published dataset for REE concentrations in terrestrial and marine vertebrates (other than humans^50^). Interestingly, ∑REE concentrations in liver were consistently higher (approx. 4 - 200 times) than in muscle for all vertebrates (Fig. 2, right panel). These differences were statistically significant for willow ptarmigan, ringed seal and brook trout (lakes only) (Welch’s ANOVA, F = 43.5, p < 0.001). Other comparisons were likely not significant due to small sample size and high inter-group variability. Regression analysis comparing muscle and liver ∑REE concentrations of all vertebrates showed only a weak correlation, with very low explanatory power (N=108, 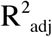 = 0.04, p = 0.03). For brook trout, however, muscle and liver concentrations showed a weak positive correlation (N=59, 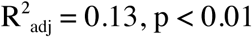).

Determining the primary organs where REEs accumulate is important for understanding the potential modes of toxic action of these contaminants of emerging concern. Previous studies on aquatic vertebrates have similarly shown that REEs are more concentrated in internal organs (liver, kidney, intestine, gills) compared to muscle.^3, 27, 44, 45^ For a marine squid, however, REE concentrations did not significantly differ between organs and muscle.^44^ There is currently little available information on the cellular mechanisms of tissue-specific REE bioaccumulation. Studies on humans have shown that our livers are often enriched in REEs (associated with proteins in intracellular complexes)^51^ and that REEs have a strong affinity for the mineral and organic components of the skeleton.^52^ Molecular levels studies are clearly needed to identify common modes of action and the biochemical effects of REEs. Future research should evaluate REE concentrations in internal organs and/or whole organism concentrations (including bones) because muscle concentrations may not provide accurate estimates of REE exposure.

### REE Anomalies and ∂^13^C

Bioaccumulation patterns appeared relatively uniform across the REE series in our dataset. However, the normalisation of individual REE concentrations to a shale standard (PAAS) revealed changes in the relative abundance of two redox sensitive elements (Ce, Eu). Whereas invertebrates displayed little deviation from a horizontal line, strong positive Eu anomalies (∂Eu) were found for all vascular plants (median of 18, range 2.4 - 48) (Fig. 4). Positive anomalies (values > 1) indicate increased uptake of the element relative to other REEs. Strong positive ∂Eu were also noted for moss and lichen samples by a previous study in the Canadian Arctic.^47^ Possible explanations for positive ∂Eu include the reduction of Eu^3+^ to the more mobile Eu^2+^ under anoxic conditions^5^ or the preferential transport of Eu into biota due to similarities between Eu^3+^ and Ca^2+^.^53^ Normalised data for vertebrate muscle tissues showed a lot of scatter and should be interpreted with care because concentrations were highly variable and close to detection limits. However, normalised liver concentrations showed a clear downward slope with LREE enriched relative to HREE (Fig. 4). This trend has also been noted in human organs (aorta, liver and bone) but the mechanisms driving this pattern remain uncertain.^51^

**FIGURE 4:**
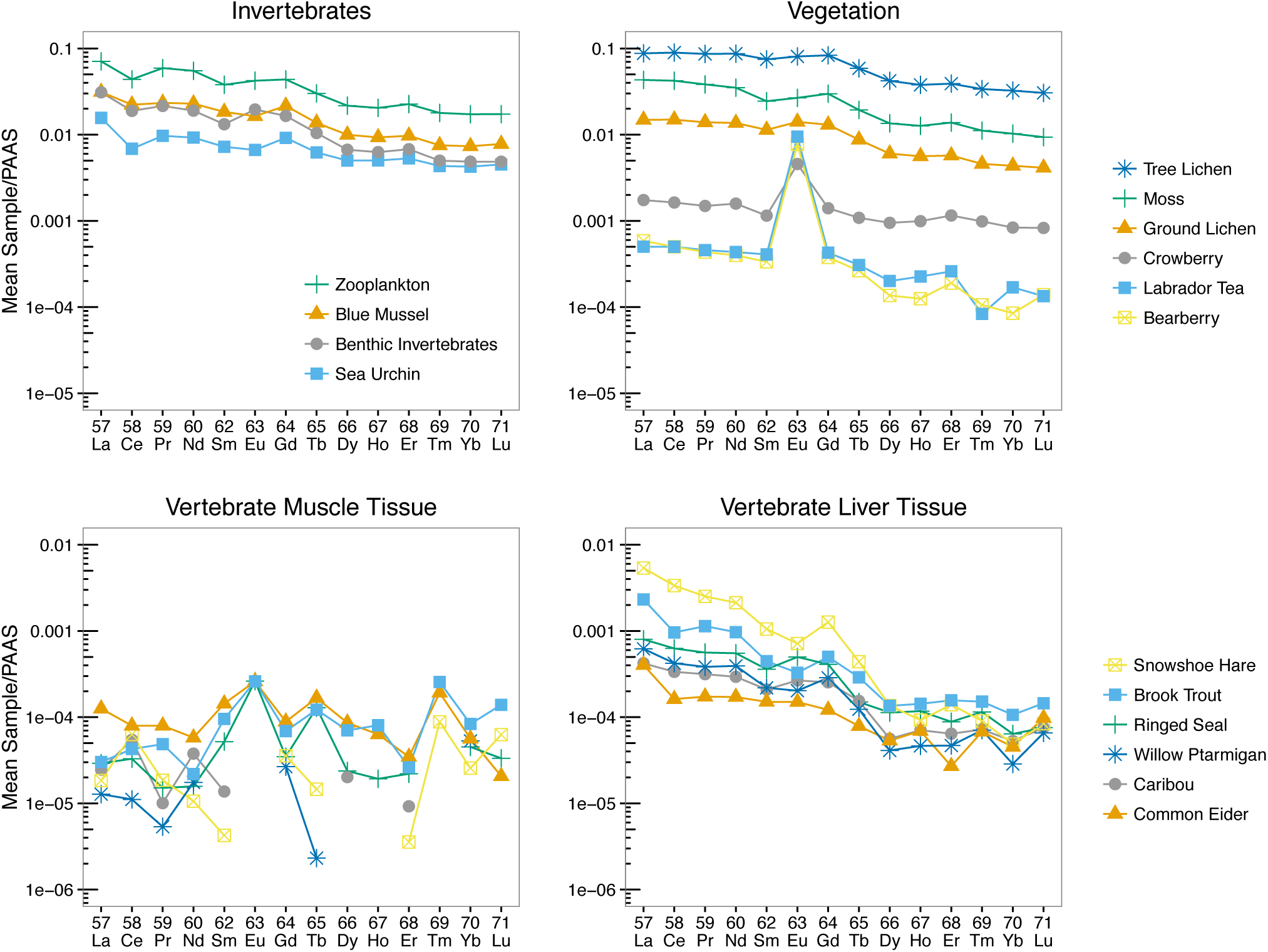
PAAS-normalized REE concentrations (geometric means, log nmolg^-1^) versus atomic number for biotic components from all ecosystems including invertebrates, vegetation, vertebrate muscle and vertebrate liver tissues. Points show element means for each taxonomic group and samples below detection limits were excluded from the figure (e.g. muscle tissues).

Interesting trends were found between carbon stable isotope ratios (∂^13^C, ‰) and Eu, Ce anomalies for some taxonomic groups. ∂^13^C and positive ∂Eu values (log_10_ -scaled) were negatively correlated for vegetation (N=15, 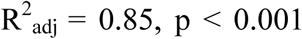) and benthic invertebrates (N=17, 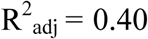) (Fig. S2). For vegetation, this shows that higher Eu accumulation occurs in C3 plants with low ∂^13^C values (around −30 ‰). For freshwater invertebrates, this shows that Eu accumulation increases in benthos feeding on a more pelagic source of carbon. A significant positive correlation was also found for ∂^13^C and negative ∂Ce values in brook trout liver (N=60, 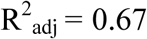, *p* < 0.001) (Fig. S3). Thus, lower Ce bioaccumulation occurred in fish feeding on plankton carbon over benthic carbon. Fish feeding more on benthic carbon had ∂Ce close to 1, indicating no significant anomaly. Correlations with ∂^13^C indicate that individual element anomalies are related to exposure routes, yet may also reflect internal physiological processes. Overall, bioaccumulated REEs behaved as a coherent group, however species- and tissue-specific anomalies occurred for some elements.

### Size and Sex-Dependent REE Bioaccumulation

For ringed seal, no evidence was found for an effect of sex (fixed effect) or year of collection (random effect) on liver REE concentrations. Seal age was not measured. A low sample size and unbalanced design may contribute to the lack of an effect in this model because a small number of female seals were sampled (N = 6). The best fit model was a simple linear regression with seal length as a significant predictor of liver REE concentration (logREE ∼ Length, 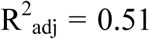, *p* < 0.001) (Fig. 5). Seal girth (axial, total) was highly collinear with seal length (r = 0.90) and was also positively correlated with liver REE concentration (logREE ∼ Girth, 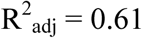, p < 0.001).

**FIGURE 5:**
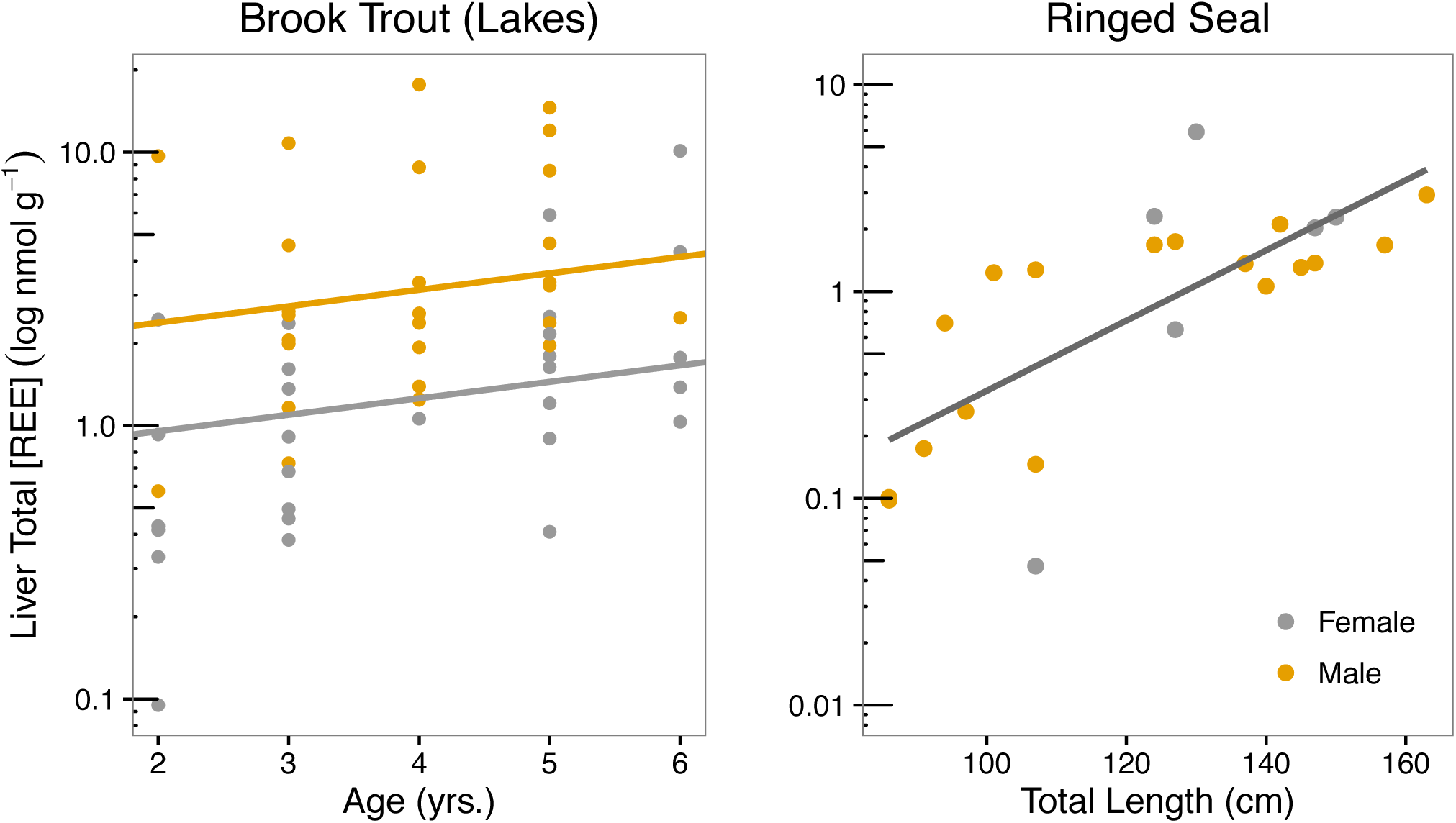
Relationships between sex and size (age or length) of brook trout and ringed seal and liver REE concentrations (log-scaled nmolg^-1^). Linear mixed effects models (LMM) indicate that sex (p < 0.001) and age (p = 0.03) were significant predictors of liver REE concentration for brook trout (N=58, sex coefficient ± SE: *female* = −0.14 ± 0.17, t value = −0.82; *male* = 0.26 ± 0.06, t value = 6.34; age coefficient (coef) ± SE: age = 0.06 ± 0.03, t value = 2.28). No significant effect of length was found for liver REE concentration in brook trout. Simple linear regression indicates that seal length (cm) was a significant predictor of liver REE concentration for ringed seal (N=23, p < 0.001).

For brook trout, there was a statistically significant relationship between age, sex and liver REE concentration (Fig. 5). The best fit LMM included the fixed effects (sex, age) and a random intercept (lake ID) (logREE ∼ Sex + Age + (1|Lake)). Fixed effects explained 25.0% of the variation in liver REE concentration (marginal R^2^) and the full model explained 72.3% (conditional R^2^); thus 47.3% of the variance was associated with the random effect, i.e. habitat or lake. The random intercept indicated that the influence of sex and age on liver REE concentration varied among fish from different lakes. There was a weak positive correlation between fish age and REEs, with REE levels increasing at approximately 1.15 nmolg^-1^ per year in fish livers. On average, male fish had liver concentrations 2.5 times higher than female fish (p < 0.001, Fig. 5). No significant overall effect of length on REE concentration was found for fish liver in this dataset, although fish length and age were only weakly related across lakes (N = 58, r = 0.31, p = 0.02). Overall, the model suggests that sex and age influenced REE bioaccumulation in brook trout but that site-specific exposure was more important.

Metal bioaccumulation occurs due to high rates of uptake from different sources (water, food or air) coupled with slow rates of elimination.^54^ These results indicate that REE bioaccumulation is greater than elimination over time for ringed seal and brook trout. Similar results were also reported by Mayfield and Fairweather for rainbow trout (also in the family *Salmoninae*) with REE concentrations increased weakly (and not always significantly) with fish age, size and weight.^27^ However, the authors also found that sucker species showed significant negative correlations with age, size or length, indicating that REE bioaccumulation patterns vary with species. An effect of sex was found only for the brook trout. Since no significant differences in length or weight were observed between male and female fish, lower liver REE concentrations in female fish could be due to metal depuration during egg production, differences in foraging behaviour, or dimorphism in liver function (but not size dimorphism). ^55^

## CONCLUSIONS

This study greatly improves our understanding of REE bioaccumulation and trophic transfer in remote Arctic ecosystems. The bioaccumulation of REEs showed a predictable and coherent pattern of log-linear decrease with atomic number for most tissues. The normalization of individual REE concentrations revealed species- and tissue-specific anomalies for the redox-sensitive elements (Ce, Eu). This study also showed that REE bioaccumulate in a wide variety of biota from marine, freshwater and terrestrial ecosystems and that primary producers/consumers are good bio-indicators of REE pollution in the environment because of their higher concentrations. REE levels decreased with trophic position, which indicated limited potential for biomagnification. Low levels of REEs in vertebrate muscles indicate that consumption of these tissues is unlikely to be an important exposure route for humans in northern regions unaffected by mining activities. Future research should focus on REE concentrations in internal organs and/or whole organism concentrations because low muscle REE concentrations may not provide accurate estimates of environmental exposure to total REEs. The findings on REE behaviour and bioaccumulation patterns from this study provide critical new information for assessing the potential toxicity pathways of REEs in vulnerable northern environments.

## Author Contributions

JH, RM and JC coordinated the collection of field data. Sampling was designed by JH, RM, and JC. Data and statistical analysis was done by GAM. GAM, JC and MA prepared the manuscript. All authors have given approval to the final version of the manuscript.

## Acknowledgements

This study is an example of fruitful research that can be completed through cooperation between the scientific and indigenous communities in the Arctic. Special thanks to all the hunters of Kuujjuarapik-Whapmagoostui: Jimmy Paul Angatookalook, Charlie Angatookalook, Michael Angatookalook, Daniel Audla, Joanna Fleming, Jordan Kroonenberg, Willie Novalinga, Simionie Papayluk, Eddy Tookoo, Samson Tooktoo, Vincent Tooktoo, and Alec Tuckatuck. The authors would also like to thank Tania Perron and Dominic Bélanger for their help in the field and in the lab. Thanks to Catherine Girard, Meredith Claydon, Emmanuelle Chrétien and Zofia Ecaterina Taranu for help with statistical analyses and with R.

